# Genetic evidence for a large overlap and potential bidirectional causal effects between resilience and well-being

**DOI:** 10.1101/2020.11.03.366492

**Authors:** L.P. de Vries, B.M.L. Baselmans, J.J. Luykx, E.L. de Zeeuw, C. Minică, E.J.C. de Geus, C.H. Vinkers, M. Bartels

## Abstract

Resilience and well-being are strongly related. People with higher levels of well-being are more resilient after stressful life events or trauma and vice versa. Less is known about the underlying sources of overlap and causality between the constructs. In a sample of 11.304 twins and 2.572 siblings from the Netherlands Twin Register, we investigated the overlap and possible direction of causation between resilience (i.e. the absence of psychiatric symptoms despite negative life events) and well-being (i.e. satisfaction with life) using polygenic score (PGS) prediction, twin-sibling modelling, and the Mendelian Randomization Direction of Causality (MR-DoC) model. Longitudinal twin-sibling models showed significant phenotypic correlations between resilience and well-being (.41/.51 at time 1 and 2). Well-being PGS were predictive for both well-being and resilience, indicating that genetic factors influencing well-being also predict resilience. Twin-sibling modeling confirmed this genetic correlation (.71) and showed a strong environmental correlation (.93). In line with causality, both genetic (51%) and environmental (49%) factors contributed significantly to the covariance between resilience and well-being. Furthermore, the results of within-subject and MZ twin differences analyses were in line with bidirectional causality. Additionally, we used the MR-DoC model combining both molecular and twin data to test causality, while correcting for pleiotropy. We confirmed the causal effect from well-being to resilience, with the direct effect of well-being explaining 11% (T1) and 20% (T2) of the variance in resilience. Data limitations prevented us to test the directional effect from resilience to well-being with the MR-DoC model. To conclude, we showed a strong relation between well-being and resilience. A first attempt to quantify the direction of this relationship points towards a bidirectional causal effect. If replicated, the potential mutual effects can have implications for interventions to lower psychopathology vulnerability, as resilience and well-being are both negatively related to psychopathology.

## 1. Introduction

In life, everyone is exposed to multiple personal adverse or stressful life events, such as a traffic accident or the death of a loved one. These adverse life event, but also events like terroristic attacks or worldwide crises, can lead to stress and trauma. There are individual differences in the responses of people to (major) life stressors and potential trauma (Luthar et al., 2000; Werner & Smith, 2001), and resilience is found to be the most common response according to a recent review of 54 studies (Galatzer-Levy et al., 2018). Resilience can be defined as the process of quickly recovering after the experience of stress or trauma (Charney, 2004; Choi et al., 2019; Connor & Davidson, 2003; Galatzer-Levy et al., 2018; Kalisch et al., 2017; Luthar et al., 2000). Resilient people adapt relatively quickly, after some time their well-being levels are back to baseline (see the review of Galatzer-Levy et al., 2018). Less resilient people do not cope well in response to stress and experience chronic or long-term adverse effects, leading to the development of psychopathology (e.g. depression) (Bonanno, 2004; Bonanno et al., 2011; Galatzer-Levy et al., 2018; Galatzer-Levy & Bonanno, 2012; Kendler et al., 2000; Pietrzak et al., 2014).

For example, in response to the COVID-19 pandemic, only 13.6% of a USA representative sample (N=1468) showed psychological distress (McGinty et al., 2020). While this level of distress is higher than the 3.9% found in the same sample in 2018, 86% of the sample seemed to be resilient, as their distress did not increase. As another example, Bonanno et al. (2006) reported widespread resilience in a large sample (N=2752) that was exposed to the September 11^th^ attacks in New York. Across all participants, 65.1% could be classified as resilient. When investigating subgroups, even in the group participants that was in the World Trade Center building at the time of the attack, more than half of the sample (53.5%) was resilient.

One of the correlates of resilience that is often suggested to play a role in bouncing back to normal is well-being. Well-being can be defined in multiple ways and a distinction between subjective and psychological well-being can be made. The subjective well-being theory originates from the hedonistic philosophy of well-being, whereas psychological well-being emerged from eudaimonic philosophy (Ryan & Deci, 2001; van de Weijer et al., 2018). Subjective well-being is characterized by high levels of positive affect and a high subjective evaluation of life satisfaction (Diener et al., 2018), whereas psychological well-being refers to thriving, positive functioning, and judgments about the meaning and purpose of an individual’s life (Ryff, 1989). It has repeatedly been found that well-being plays a preventive role in psychopathology and is important to overall physical and mental health (Diener et al., 2017; Greenspoon & Saklofske, 2001; Howell et al., 2007). Well-being associates positively with longevity and health (James et al., 2019; Kim et al., 2019; Steptoe, 2019; Zaninotto & Steptoe, 2019), satisfaction with marital relationships, productivity at work, prosocial behavior and educational achievement (Chapman & Guven, 2016; Lyubomirsky et al., 2005; Maccagnan et al., 2019; Oswald et al., 2015).

A strong positive correlation (around 0.5) between resilience and well-being has been found as well (e.g. Bajaj & Pande, 2016; Hu, Zhang, & Wang, 2015; Liu, Wang, & Li, 2012; Satici, 2016). That is, people with a higher well-being show more resilience and, vice versa, people who are more resilient show higher well-being (Cohn et al., 2009; Ong et al., 2006). For example, women reporting higher life satisfaction are more likely to be resilient after their spouse passes away (O’Rourke, 2004; Rossi et al., 2007). Fredrickson et al. (2003) suggests that positive emotions are the active elements in resilience. In times of crisis, the presence of positive emotions buffers against depression and adverse outcomes. Conversely, Connor and Davidson (2003) suggest resilience is a protective factor in facing negative emotions after adverse events and therefore protects people’s well-being. These studies, though, are correlational, and causal interpretation is hard. It is theoretically plausible to expect a bidirectional relation between resilience and well-being. When people are able to cope with life stressors and adapt well to adversity (resilience), they feel better and evaluate their life positively (well-being) compared to people that cannot cope well. In turn, positive emotions and higher levels of well-being improve the ability to respond adaptively to life events, i.e. resilience.

Alternatively, underlying genetic or environmental confounders might induce the association between well-being and resilience - without a direct causal effect in either direction. The genetic and environmental factors contributing to both phenotypes have been investigated before. Two meta-analyses summarized all studies applying the twin model to well-being and found a meta-analytic heritability (i.e. the contribution of genetic factors to the variance) of 36% (CI: 34-38%) and 40% (CI: 38-43%) for well-being based on all measures (Bartels, 2015; Nes & Røysamb, 2015) and 32% (CI: 29-35%) for satisfaction with life (Bartels, 2015). The remaining variance was explained by unique environmental influences. Causes of individual differences in resilience have also been investigated, although less frequent and with substantial variation in operationalization, sample size, and sample composition (e.g. population vs military sample). In two studies based on military samples, the heritability estimates of self-reported resilience were 25% (CI: 21-30) and 55% (CI: 48-61) respectively (Long et al., 2017; Wolf et al., 2018). In adolescents and based on three raters (father, mother, self) a common resilience factor (excluding rater specific error) showed heritability estimates of 78% in boys and 70% in girls (Waaktaar & Torgersen, 2012). When operationalizing resilience as the residual of positive affect after controlling for stressors or the residual of internalizing symptoms after controlling for stressful life events, heritability estimates ranged from 31% to 52% (Amstadter et al., 2014; Boardman et al., 2008). Using multiple measures to reduce measurement error, Amstadter et al. (2014) found a heritability of 50% (CI: 46-59) for the latent construct of resilience.

The above studies show that resilience and well-being are strongly related phenotypically and have a similar genetic architecture. The etiology of well-being and resilience has not been addressed in a bivariate design, to formally test the overlap in genetic and environmental factors. In order to get a better hold on the nature of the association between resilience and well-being, and possible roles for genetic confounding and (bidirectional) causal effects we took a three-step approach. (1) First, we estimated the crosssectional and longitudinal phenotypic association between well-being and resilience. (2) Next, we used genome-wide summary statistics of well-being to predict resilience and wellbeing. (3) Finally, we tried to falsify the causal hypotheses (in both directions) using various approaches: within-subject change score regression, bivariate psychometric twin-sibling modeling, cross-trait correlations of intrapair MZ twin differences, and the MR-DoC model which combines Mendelian Randomization and the Direction of Causation twin model.

## 2. Method

### 2.1. Sample

Participants were registered at the Netherlands Twin Register (NTR), established by the Department of Biological Psychology, Vrije Universiteit Amsterdam (Ligthart et al., 2019). The NTR sample is a population-wide, non-clinical sample. Every two/three years, longitudinal survey data about lifestyle, personality, psychopathology, and well-being in twins and their families are collected. The current study used data on life events, psychopathology, and well-being in adults collected in 2002-2003 (Time point 1) and 2009-2012 (Time point 2) (Ligthart et al., 2019; Willemsen et al., 2013).

Including twins and biological siblings, the total sample consisted of 14.055 participants with data on resilience and/or well-being collected in one or both waves. We excluded 177 participants with unknown zygosity and two participants with unknown sex, resulting in a final sample of 13.876 participants (11.304 twins and 2.572 siblings). The sample included 1.577 monozygotic male (MZM), 967 dizygotic male (DZM), 3.859 monozygotic female (MZF), 2.112 dizygotic female (DZF) and 2.789 dizygotic opposite-sex (DOS) twins from complete and incomplete twin pairs.

When split by time of data collection, 4.447 twins and 1.407 siblings (33.8% male, *M_age_*= 32.87, *SD_age_* = 11.41) had data at time point 1 (T1). At time point 2 (T2), data of 9.590 twins and 1.962 siblings (32.3% male, *M_age_*=31.73, *SD_age_*= 14.41) were available. Longitudinal data (both at T1 and T2) were available for 3.530 participants (2.733 twins and 797 siblings, 41.1% male, T1: *M_age_*= 33.91, T2: *M_age_*= 40.25, *SD_age_*= 11.6 (see Supplementary Table S1 for more information).

The data collection was approved and declared to be of low risk and exempt of formal medical ethical risk assessment by the METc of the Vrije Universiteit Medical Center Amsterdam (Approval: NL25220.029.08 (ref # 2008/244), 1 December 2008 and 2011/334, 12 Oct 2011; 2012/433, dd 26 Feb 2013).

### 2.2. Measures

#### 2.2.1. Well-being

Well-being was assessed with the Satisfaction with Life scale (Diener et al., 1985). The scale consists of five items with a 7-point Likert scale, ranging from *1 = strongly disagree* to *7 = strongly agree*. An example question is ‘*In most ways my life is close to ideal*’. Items were summed to calculate an individual’s final score ranging from 0 to 35, with higher scores indicating higher levels of satisfaction with life.

#### 2.2.2. Life events

The number of experienced life events was assessed with an adapted version of a Dutch life-event scale (Schokverwerkings Inventarisatie Lijst = SchIL; Van der Velden et al., 1992). At T1, 16 negative life events items were included about the experience of death of a significant other, serious disease of yourself/significant other, end of relation, traffic accident, violent or sexual assault, robbery, and getting fired. Time point 2 included 28 life events, both positive and negative events. In line with previous work, we excluded the positive life events at T2, leaving 19 items (Middeldorp et al., 2008). Possible answers were *never experienced, experienced last year (0-12 months), 1-5 year ago and >5 years ago*. The number of life events was computed by summing the experienced life event in the past 5 years. The maximum number of life events experienced is 16 at T1 and 19 at T2 (see Supplementary Table S2).

#### 2.2.3. Anxious-depressed

Anxious-depressed symptoms were assessed with the anxious-depressed subscale of the Adult Self Report of the Achenbach System of Empirically Based Assessment (ASEBA; Achenbach & Rescorla, 2003). Each item is rated from 0 = *not true*, 1 = *somewhat true*, to 2 = *very true*. An example item is *“I feel worthless”*. As T1 only included 15 (instead of 18) items of the scale, at T2s we only selected those same items and created a sum score of 15 items, with higher scores indicating higher levels of anxious-depressed behavior.

#### 2.2.4. Resilience score

Resilience is operationalized as an outcome-based measure in line with Amstadter et al. (2014) and is based on the regression of internalizing problems on the total number of stressful life events experienced (e.g. Kendler, Thornton, & Gardner, 2000; Kessler, 1997; Phillips, Carroll, & Der, 2015). Resilience is defined as the difference between the predicted level of anxious-depressed symptoms based on the number of life events and the actual level. Individuals who experience less anxious-depressed symptoms than expected based on the number of stressful events in their life can be seen as resilient.

To this end, the number of life events and the anxious-depressed scores were standardized. For both T1 and T2, the resilience score was operationalized as the residuals of the anxious-depressed sum score after the effect of the number of stressful life events had been regressed out, using Generalized estimating equation (GEE) to correct for familial relations (Minică et al., 2015). This standardized residual is our measure for resilience and used in further analyses.

#### 2.2.5. Genotype data

Genotype and phenotype information was available for 10867 NTR participants in our sample. Genotyping was done on several genotyping arrays, including the Axiom array (N=615), Affymetrix 6.0 (N=6144), Illumina Omni Express 1 M (N=181), Illumina 660 (N=1312), Illumina GSA (N=4044) and Perlegen/Affymetrix (N=1013) (see Ehli et al., 2017; Willemsen et al., 2013). Additionally, SNPs extracted from sequence data from the Netherlands reference genome project Genome of the Netherlands (GoNL) (N=267) were used (Boomsma et al., 2014; The Genome of the Netherlands Consortium, 2014).

For each platform, SNPs with a Minor Allele Frequency (MAF) <0.01 or SNPs out of Hardy–Weinberg Equilibrium (HWE) with p< 10^−5^ were removed. Also, samples were excluded if there was a mismatch in expected and genotyped sex, the genotype missing rate was above 10% or the inbreeding value (Plink F statistic) was not between −0.10 and 0.10. To control for Dutch population stratification, Principal Components Analysis (PCA) was performed and individuals with a non-Dutch ancestry based on their PCs were excluded, as described by Abdellaoui et al. (2013).

To infer the SNPs missing per platform in the combined data, the genotyped data of the different arrays were cross-platform imputed using the GoNL as a reference panel (Boomsma et al., 2014; Consortium et al., 2014; Fedko et al., 2015). SNPs were removed if alleles mismatched with the reference panel, were out of HWE with p < 10^−5^, the Mendelian error rate was larger than the mean + 3 SD, or the imputation quality (R2) was below 0.90. The SNPs in the final cross-platform imputed dataset were aligned similarly to the 1000 Genomes Phase 3 v5 reference panel, and uploaded to the Michigan Imputation Server. Here, the data were phased and imputed to this 1000 genomes panel using Shapeit and Minimac3 respectively. The data were again filtered for SNPs having MAF < 0.01, HWE p < 10^−5^, alleles not being A, C, G, or T, and a call rate of 99% after all Mendel errors were removed. A random selection of 2500 second degree unrelated people were taken from this dataset. Using the summary statistics, this unrelated set and LDpred, the beta’s were corrected. As described in more detail below, polygenic scores were constructed on the relevant 10867 individuals.

### 2.3. Analyses

#### 2.3.1. Part 1. Demographics and phenotypic correlations

First, we applied a saturated twin-sibling model in OpenMx (Boker et al., 2011) including the resilience and well-being scores at both time points to test the equality of means and variances in twins and siblings, and sex differences in the means of resilience and wellbeing. Furthermore, the cross-sectional and longitudinal phenotypic correlations and twin and twin-sibling correlations within and across traits were estimated.

#### 2.3.2. Part 2. Genetic Prediction

To investigate the molecular genetic overlap between resilience and well-being, we used summary statistics of two recent genome-wide association studies (GWASs) on resilience and well-being to create polygenic scores (PGS). PGS are a measure of an individual’s genetic probability to develop a certain disorder or have a certain trait (Wray et al., 2007). Using GWAS summary statistics, the PGS of a phenotype can be calculated in an independent sample by summing all genotype scores (at individual single-nucleotide polymorphisms) for a person after weighting them by their estimated effect size. The PGS of the phenotype can be used to test the predictive value towards another trait, or to investigate the shared genetic etiology between traits (Purcell et al., 2009).

We used PGS for resilience and well-being to investigate if and to what extent the genetic risk for well-being is a predictor for resilience and vice versa. For well-being, the polygenic scores from the most recent GWAS summary statistics for the well-being spectrum, leaving out NTR, were used (Baselmans, Jansen, et al., 2019). Using the summary statistics of the only GWAS to date on self-assessed resilience based on a sample of 11,492 army soldiers (Stein et al., 2019), we created PGS scores for resilience in the NTR sample.

The polygenic scores were computed using LDpred (Vilhjálmsson et al., 2015). LDpred takes into account linkage disequilibrium (LD) among SNPs in creating the polygenic risk scores. We calculated the mean causal effect size of each marker using the SNP effect sizes from the resilience and well-being summary statistics. The LD structure from a reference set specific for the NTR based on 1000 Genomes phase 1 genotypes (1000 Genomes Project Consortium, 2015) was used to calculate polygenic scores in the target sample, in this case the NTR sample. In order to avoid an over-estimation of the association between the polygenic scores and phenotypes, summary statistics for the well-being GWAS in the discovery set were re-computed, excluding NTR subjects. Polygenic scores were calculated with the fractions of causal genetic variants (the fraction of markers with non-zero effects) set to 1, 0.5, 0.3, 0.2, 0.1,0.05, and 0.01 to test which fraction suited the data best. We restricted analyses to common variants, using a SNP inclusion criterion of minor allele frequency (MAF) > 5%.

Using GEE to correct for familial relations, we regressed the created PGS of resilience on well-being and vice versa and included age, age^2^, sex, the genotyping array, and the first ten genomic principal components (PCs) as covariates. A significant association indicates that the genetic risk for resilience predicts well-being or vice versa. To correct for multiple testing, we used a Bonferroni corrected threshold of 0.001 for significance.

#### 2.3.3. Part 3. Causality

Next, we investigated the possible direction of causation between resilience and wellbeing. Under the causal hypothesis, several predictions in cross-sectional and longitudinal data can be made (Bartels et al., 2012; De Moor et al., 2008), that are specified in in section 2.3.3.1 until 2.3.3.4. Importantly, with these test we will not be able to confirm causality, but we are able to falsify the causal hypothesis.

##### 2.3.3.1. Within-subject change scores

First, we used regression of the within-subject changes in well-being and resilience over time. If there is a causal relation from resilience to well-being, within-subject changes in resilience over time (T2 – T1) should predict parallel changes in well-being over time. Under the causal hypothesis, increases in resilience over time would result in increase in well-being. The absence of a correlation of change scores over time would reject the causal hypothesis, whereas the presence of a correlation is in line with causality (Bartels et al., 2012; De Moor et al., 2008). Using GEE to correct for relatedness, regression analyses were performed to predict within-subject changes in well-being by within-subject changes in resilience over time. In reverse, if there is a causal relation from well-being to resilience, within-subject changes from T1 to T2 in well-being should predict parallel changes in resilience over time. These regression analyses exclude confounding by genetic factors, since the genotype within a subject does not change.

##### 2.3.3.2. Bivariate twin-sibling models

Another prediction under the causal hypothesis is that if resilience is causally related to well-being, all genetic and environmental factors that influence resilience will also, through the causal chain, influence well-being (De Moor et al., 2008).

To test the significance of genetic and environmental correlations between resilience and well-being, we used twin-sibling models. Bivariate twin-sibling models use the difference in genetic overlap between monozygotic (MZ) twins and dizygotic twins (DZ) to estimate the underlying sources of phenotypic variance of two traits and their phenotypic correlation. In addition, these model results can be used to calculate genetic and environmental correlations (de Vries et al., BG). MZ twin pairs are genetically identical, whereas DZ twin pairs share on average half of their segregating genes. Based on this difference, the observed phenotypic variance and covariance between traits can be decomposed into genetic and environmental variance components Additive genetic variance (A) is the variance explained by the independent allele effects on the phenotype. Non-additive genetic variance (D) refers to interactions between alleles at the same locus (dominance) or between alleles at different loci (epistasis). Environmental variance includes a shared environmental variance component (C) (shared by family members) and a non-shared component, the unique environment, also including measurement error (E). The effects of C and D cannot be estimated simultaneously for identification reasons and a choice between an ACE or ADE model is made based on twin correlations. The power of the classical twin design increases by adding non-twin siblings of twin pairs. These non-twin sibling share on average half of their segregating genes with other siblings (including the twins) and can be treated as DZ twins in the models (Posthuma & Boomsma, 2000).

Using the log-likelihood ratio test (LRT), the full ACE/ADE models were compared to nested submodels. The difference in minus two times the log-likelihood (−2LL) between two nested models has a χ^2^ distribution with the degrees of freedom (df) equaling the difference in df between two models. If a p-value from the χ2-test was higher than the alpha of 0.001 (corrected for multiple testing), the constrained and more parsimonious model fit was not significantly worse than the fit of the more complex model. The distribution of resilience and well-being scores were moderately skewed, but showed a bell-shaped curve and were therefore analyzed as continuous variables. Furthermore, whereas the skewed data might bias the parameter estimates, transformations do not remove the known and small bias (underestimation of the shared environmental effect, and overestimation of the unique environmental effect (Derks et al., 2004)).

As we have data on resilience and well-being at two time points, we modelled the variance of the underlying phenotypes in a bivariate psychometric model with repeated measures. The resilience and well-being scores at T1 and T2 can be seen as an index of the true measure including measurement error (Amstadter et al., 2014). For both resilience and well-being, the variance was split into a common (latent or stable) part and two uncorrelated (time-specific) parts. Next, both the common and time-specific parts of the variance were decomposed in variance explained by A, C/D, and E. The variance of the latent factors includes less measurement error, therefore this results in more reliable estimates of the genetic and environmental effects (see supplementary Figure S1 for the model) and is more comparable to the earlier work by Amstadter et al. (2014).

To investigate the overlap and genetic architecture of the latent factors of resilience and well-being, we estimated genetic and environmental contributions to the variance and covariance of the latent factors. Furthermore, the genetic and environmental correlations are calculated. In this model, we first tested for quantitative sex differences (i.e. if the estimates of the genetic contribution in males and females are similar) by constraining the estimates of A, C/D and E to be equal in males and females. Next, we estimated the contribution of the variance components A and C/D to the total variance and covariance of the phenotypes. We did not test for qualitative sex differences, as modelling sex specific genes in multivariate models has inherent limitations (Neale et al., 2006) and no qualitative sex effects in well-being are expected (Stubbe et al., 2005).

If resilience and well-being are causally related, genetic and environmental factors influencing individual differences in one trait will, through the causal chain, also influence individual differences in the other trait. To test this causal effect hypothesis, we tested the genetic or environmental correlation between the latent traits in the bivariate model. Both the genetic and environmental correlation should be significant if there is causality. A significant genetic correlation but a non-significant environmental correlation falsifies the causal hypothesis and a common genetic factor is then more likely to underlie the association between resilience and well-being.

##### 2.3.3.3. Longitudinal twin-sibling model

In a similar way, we can use the longitudinal data of resilience and well-being in a bivariate model (De Moor et al., 2008). If resilience causes higher levels of well-being, there should be a significant longitudinal association between resilience at baseline and well-being at follow-up. Similarly, if wellbeing causes higher levels of resilience, there should be a significant longitudinal association between well-being at T1 and resilience at T2. These phenotypic associations should be paired to significant genetic and environmental correlations. This was tested in a bivariate genetic model by testing the significance of the genetic and environmental correlations between resilience at baseline (T1) and well-being at a later time point (T2) and vice versa (well-being at T1 and resilience at T2).

##### 2.3.3.4. MZ twin difference model

Another prediction made by the causal hypothesis is that the within–twin pair differences of genetically identical (MZ) twins in resilience should be associated with within–twin pair differences in well-being. We applied the monozygotic within-twin pair differences method. If there is a causal relation, the MZ twin differences (Resilience _twin 1_ – Resilience _twin 2_) in resilience should be associated with within-twin pair differences in well-being (Well-being _twin 1_ – Well-being _twin 2_) and vice versa. The twin who is more resilient should have a higher well-being score than the co-twin who is less resilient. At both time points, we regressed the MZ intra pair differences in resilience on the difference in well-being and vice versa. Since monozygotic twins are genetically identical, this test excludes confounding by genetic and shared environmental factors. However, if there is an association, also other factors in the non-shared environment of the twins can underlie this association.

Additionally, we tested whether longitudinal MZ twin intrapair differences (i.e. differences in individuals’ changes) in resilience over time are associated with intrapair differences in individuals’ changes in well-being over time and vice versa. Again, significant associations are in line with a causal hypothesis. The twin who has a larger increase in resilience should have a larger increase in well-being than the co-twin who showed less increase in resilience. To test this association, we created within-individual change scores of resilience and well-being and the difference between these change scores of MZ twin pairs. We regressed the MZ intra pair differences in resilience on the difference in well-being and vice versa.

##### 2.3.3.5. MR-DoC model

To explicitly test causality, allowing for coexisting genetic confounding, we leveraged the unique database of the Netherlands Twin Register and applied the Mendelian Randomization-Direction of Causation (MR-DoC) model. The MR-DoC model uses twin data and polygenic scores, combining the strengths of Mendelian Randomization and the Direction of Causation twin model (Minică et al., 2018). In Figure 1 the MR-DoC model is presented. The black box indicates the DoC model part and the grey box indicates the Mendelian Randomization part.

**Figure 1.**
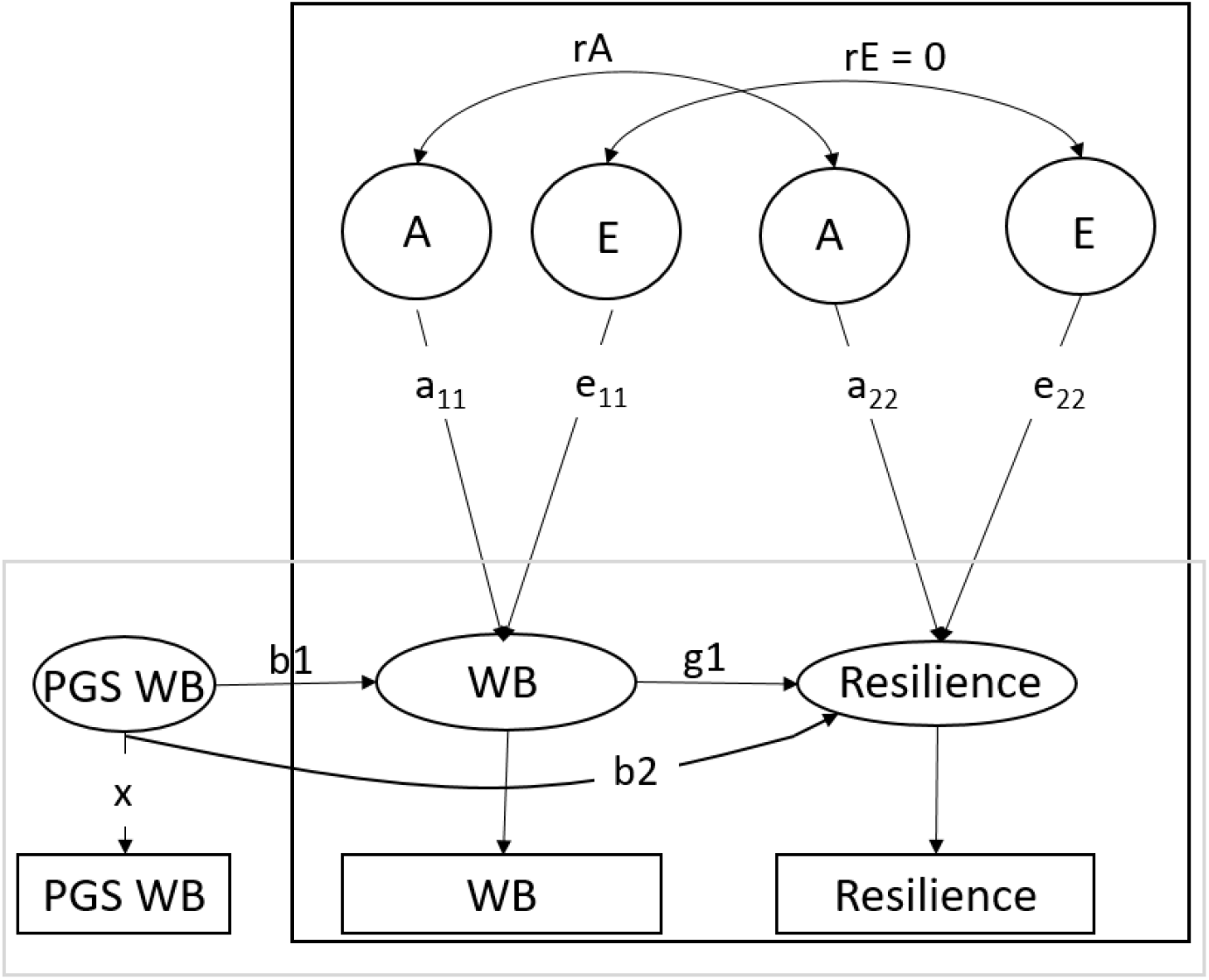
The MR-DoC model. The black box indicates the Direction of Causation model part. The grey box indicates the Mendelian Randomization part. Path g1 indicates the causal effect, path b1 indicates the PGS effect on well-being and path b2 reflects the pleiotropy between well-being and resilience. WB= well-being, A= common additive genetic effect, E= common unique environmental effects, rA= additive genetic correlation, rE= environmental correlation.

In the traditional direction of causation twin (DoC) models (Duffy & Martin, 1994; Heath et al., 1993), the covariance between traits and across twins (i.e. the cross-twin cross-trait covariance) can be used to test a causal effect from one trait on the other. The DoC model tests whether the cross-twin cross trait correlations in MZ and DZ twins reflect a unidirectional or bidirectional causal effect or a common genetic factor driving the association between the traits (i.e. significance of path g1 in Figure 1). However, to be able to distinguish between a causal effect and a common genetic factor in a DoC model, the traits do need to differ in their heritability or the sources of variance (i.e. ACE for trait 1 and AE for trait 2).

In Mendelian Randomization, genetic variants are used to test causal relationships between an exposure variable and outcome (Smith & Ebrahim, 2003). The genetic variants used to probe causal hypotheses are assumed to be: (a) well associated with the exposure variable; (b) not associated with confounders of the exposure-outcome relationship, and (c) associated with the outcome only through exposure (i.e. absence of horizontal pleiotropy). PGS can be used as strong genetic variables, but horizontal pleiotropy (assumption c) is likely to occur with complex traits (Bulik-Sullivan et al., 2015).

Pleiotropy can be divided into direct pleiotropy (i.e. a gene has a direct causal effect on multiple phenotypes, indicated by path b2 in Figure 1) and indirect pleiotropy. Indirect pleiotropy is when a gene has a causal effect on a phenotype, which in turn causally influences another phenotype (path b1*g1), indicating a causal effect.

By combining MR with twin models, the MR-DoC model can estimate both the causal effect and the amount of pleiotropy using the polygenic scores and the covariance structure between the traits, even when the traits have a similar heritability or underlying sources of variance (i.e. both AE traits). Using the cross-twin cross-trait correlations of MZ and DZ twins (like in the standard DoC model), the causal path (g1) between the traits can be estimated. At the same time, using the polygenic score, the MR part of the model normally estimates the causal effect from b1 and the observed covariance between PRS and the outcome trait (using covariance = g1*b1), assuming pleiotropy to be absent (path b2=0). As g1 is estimated from the twin DoC part, combining the covariance structure and effect of the PGS on the outcome trait, pleiotropy (path b2) can now be directly estimated. Moreover, when estimating the causal effect in the twin DoC part, pleiotropy is accounted for (for more details and simulations, see Minică et al. (2018)). Empirical analysis of height and educational attainment indicated that the test of causality conducted with MR-DoC is relatively robust to assumption violation, such as the presence of pleiotropy or assortative mating (Minică et al., 2020).

When traits have the same genetic architecture (e.g. both AE models), as is often the case, but problematic for the DoC part of the model (see Duffy & Martin, 1994), the environmental correlation between traits has to be constrained to zero for identification purposes.

We tested whether well-being causally affects resilience using the well-being PGS, the exposure being the well-being score and the outcome being the resilience score at time point T1 and T2 separately (see Figure 1 for the model). If the estimate for the causal effect from well-being to resilience (g1) is larger than zero, there is a causal effect from well-being to resilience. The b2 estimate reflects the pleiotropy between well-being and resilience. Based on the results, the effect size (% variance) of the directional effect can be estimated, taking into account the presence of residual genetic pleiotropy.

## 3. Results

### 3.1. Operationalization of resilience

The definition of the resilience assumes a positive association between stressful life events and anxious-depression and variability in the anxious-depressed score after stressful life events. Consistent with this definition, people differ in their response to stressful life events, i.e. the variance around the point estimate of anxious-depressed score increased when the number of life events experienced increased (see supplementary Figure S2). The number of stressful life events experienced and the anxious-depressed score were positively related at T1 (r=.11 [95% CI: .08-.13], β_gee_=11, p<.001) and T2 (r=.26 [95% CI: .24-.28], β_gee_=27, p<.001). The residuals from the GEE models were used as the measure for resilience.

### 3.2. Part 1. Demographic effects and phenotypic correlations

In a saturated twin-sibling model, the mean score for well-being could be constrained to be equal across males and females, (p=.113). For resilience, the means could not be constrained to be equal across sexes (p<.001). The resilience score for men was significantly higher compared to women, indicating that on average men are more resilient than women. The descriptives are given in Table 1.

**Table 1.** Descriptives of the measures for resilience and well-being.

|  |  | Time 1 |  |  | Time 2 |  |  |
| --- | --- | --- | --- | --- | --- | --- | --- |
|  |  | *N* | *M* | *SD* | *N* | *M* | *SD* |
|  | Anxious-depression | 5641 | 6.16 | 5.37 | 10804 | 4.99 | 5.05 |
|  | Number of life events | 5791 | 1.53 | 1.28 | 9719 | 2.19 | 1.84 |
|  | Well-being | 5790 | 26.52 | 0.07 | 11497 | 27.29 | 0.06 |
| Male | Resilience | 1894 | 2.97 | 0.21 | 2844 | 1.94 | 0.19 |
| Female | Resilience | 3681 | -2.02 | 0.16 | 6174 | -0.84 | 0.14 |

*Note:* the means and standard deviation for anxious-depression and number of life events are unstandardized. To compute the resilience score, these scores were standardized.

There is a small, but significant effect of age (T1: β=-.03, SE=.01, p<.001, T2: β=-.02, SE=.01, p<.001) and age^2^ (T1: β=-0.01, SE<.01, p<.001, T2: β<-0.01, SE<.01, p<.001) on well-being in both waves. Similarly, there is a small effect of age and age^2^ (all: β<0.01, SE<.01, p<.001, T2: β<0.01, SE<.01, p<.001) on the resilience score. This reflects a U-shaped curve for both well-being and resilience, indicating that younger and older people score higher on resilience and well-being than people in middle adulthood.

Table 2 shows the phenotypic correlations between resilience and well-being cross-sectionally and across the different time points. The cross-sectional phenotypic correlations between resilience and well-being are .46 (95% CI: .44-.48) and .51 (95% CI: .50-.52) at T1 and T2 respectively. The longitudinal phenotypic correlations are .35 (95% CI: .34-.36) for resilience at T1 and well-being at T2 and .43 (95% CI: .43-.44) for well-being at T1 and resilience at T2.

**Table 2.** Phenotypic correlations (with 95% CI) between resilience and well-being within and across time points.

|  | Well-being 1 | Well-being 2 | Resilience 1 | Resilience 2 |
| --- | --- | --- | --- | --- |
| Well-being 1 | 1 |  |  |  |
| Well-being 2 | 0.55<br>(.54-.56) | 1 |  |  |
| Resilience 1 | 0.46<br>(.44-.48) | 0.35<br>(.34-.36) | 1 |  |
| Resilience 2 | 0.43<br>(.43-.44) | 0.50<br>(.50-.52) | 0.61<br>(.60-.62) | 1 |

### 3.3. Part 2. Genetic Prediction

The polygenic score predictions of resilience and well-being using the different fractions of included SNPs (1 to 0.01) are in Supplementary Figure S3. The prediction of polygenic scores using a fraction of 0.5 are optimal, therefore we proceed with a fraction of 0.5. The GEE analyses show that the PGS of direct self-assessed resilience is not significant in predicting our indirect resilience score at T1 (p=.248) and T2 (p=.002), predicting only around 0.04-0.2% of the variance. The prediction of well-being by the resilience PGS is not significant (T1: p=.822 and T2: p=.144) and close to zero (see Figure 2, left panel).

**Figure 2.**
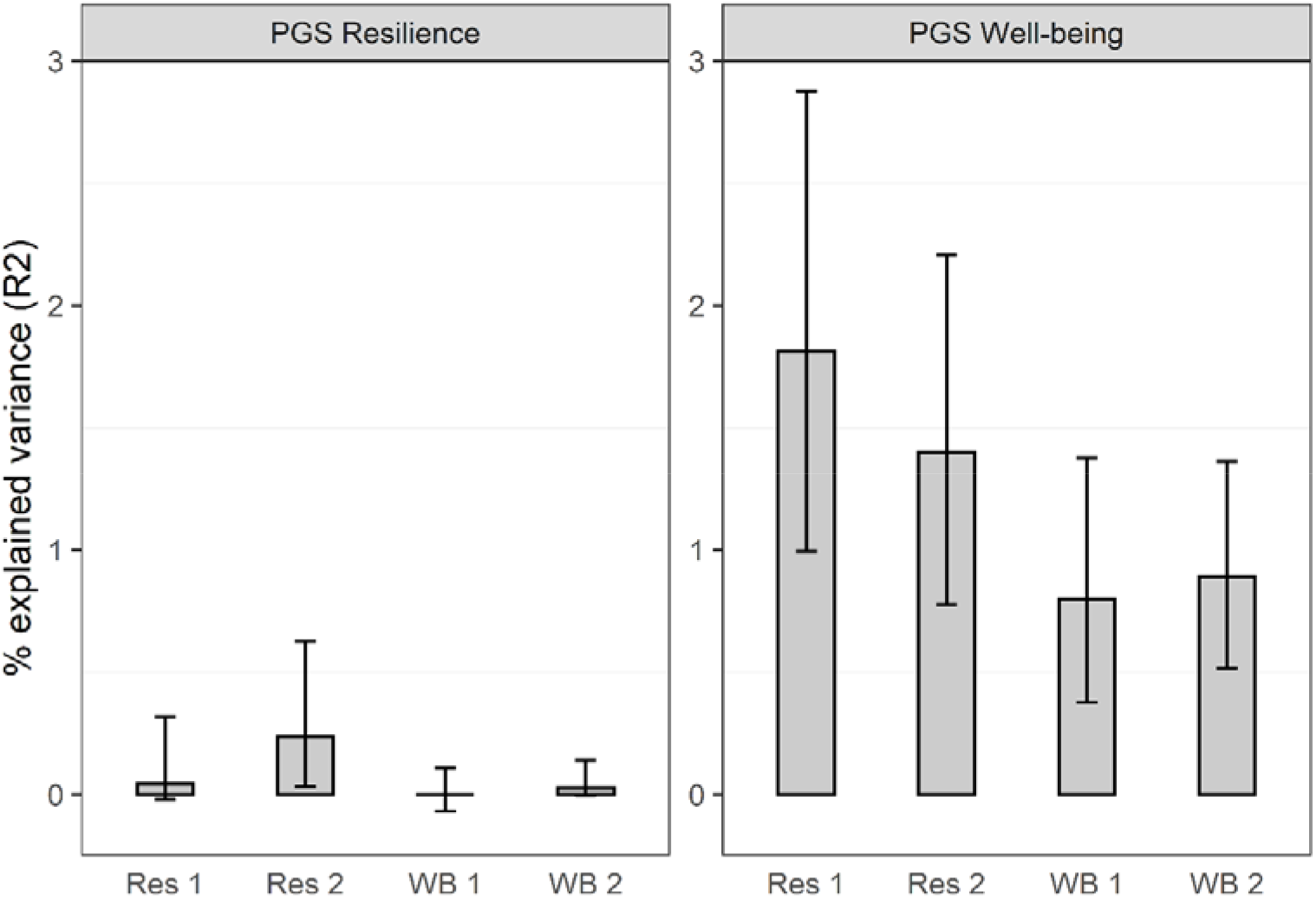
Explained variance in the phenotypic resilience and well-being scores by the polygenic scores (PGS) of resilience (left panel) and well-being (right panel). Res 1= resilience time point 1, Res 2= resilience time point 2, WB 1= well-being time point 1, WB 2= well-being time point 2.

The well-being PGS is a significant predictor for both well-being and resilience at both time points (p<.001), explaining around 0.8-0.9% of the variance in well-being and 1.4-1.8% of the variance in resilience (see Figure 2, right panel), suggesting genetic overlap between resilience and well-being.

### 3.4. Part 3. Causality

#### 3.4.1. Within-subject change scores

A change in resilience in an individual over time predicted a parallel change in well-being over time, β=.33 (95% CI: .29 -.38), SE=0.02, Z=15.67, p<.001. Similarly, within individual change in well-being predict a parallel change in resilience over time, β=.34 (95% CI: .30 -.38), SE=0.02, Z=15.95, p<.001 (see Figure 3). These results are in line with a possible causal relation between resilience and well-being, indicating that increased wellbeing can lead to increased resilience and/or vice versa, after genetic confounding is taken into account.

**Figure 3.**
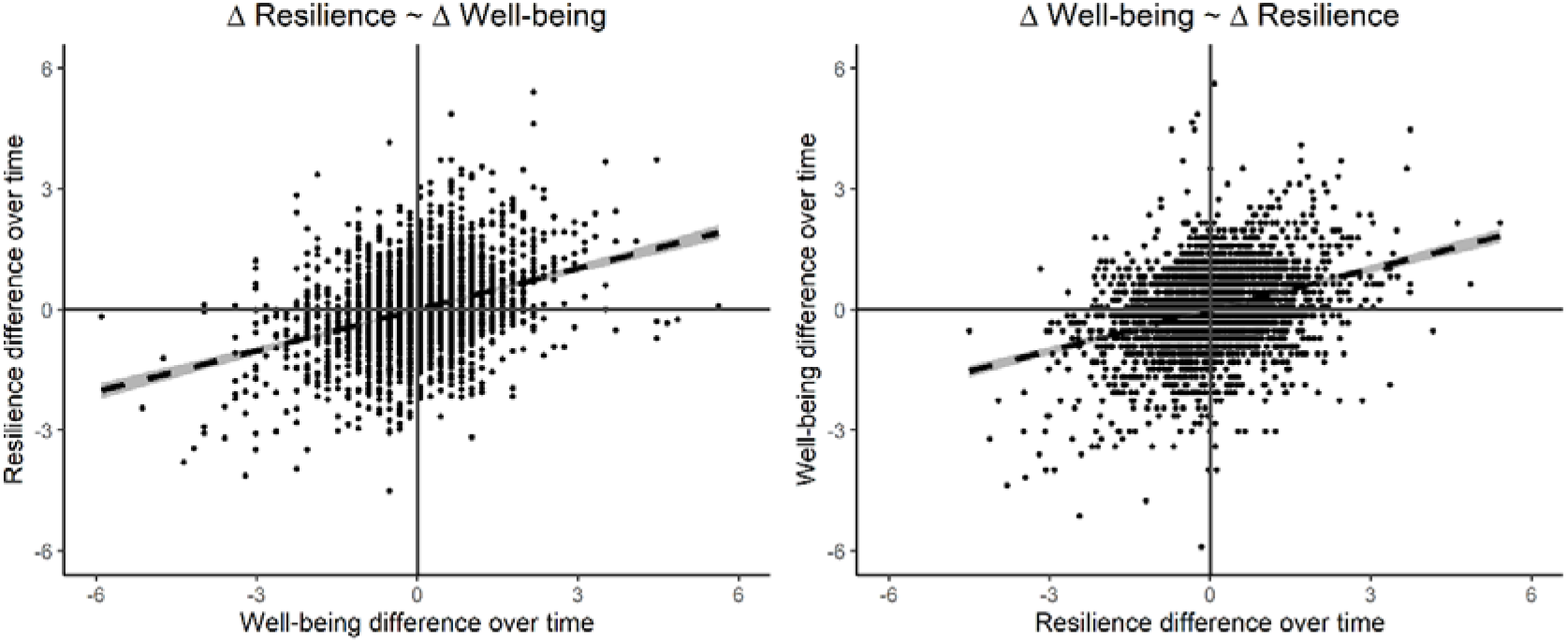
The relation between the within-subject differences over time in resilience and well-being.

#### 3.4.2. Bivariate twin-sibling models

Table 3 shows the twin and twin-sibling correlations for resilience and well-being within and cross traits and time points. Overall, the MZ correlations are more than twice the DZ/sibling correlations, suggesting dominant genetic effects besides additive genetic effects. Therefore, we continued with ADE models. Constraining all sibling correlations to the DZ correlations did not deteriorate the fit (p=.297), indicating that DZ twins do not resemble each other more than siblings (i.e. no specific twin environment).

**Table 3.**
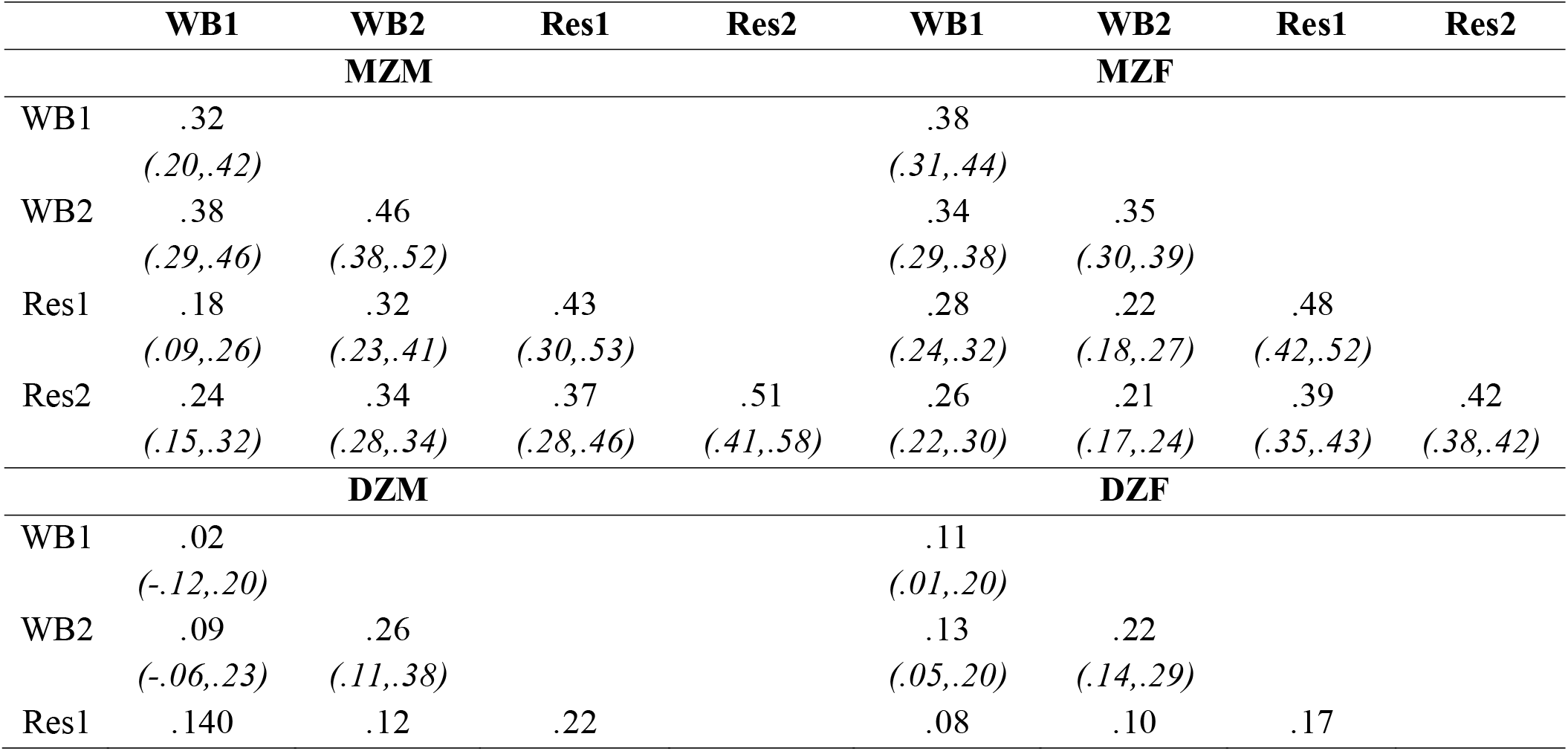
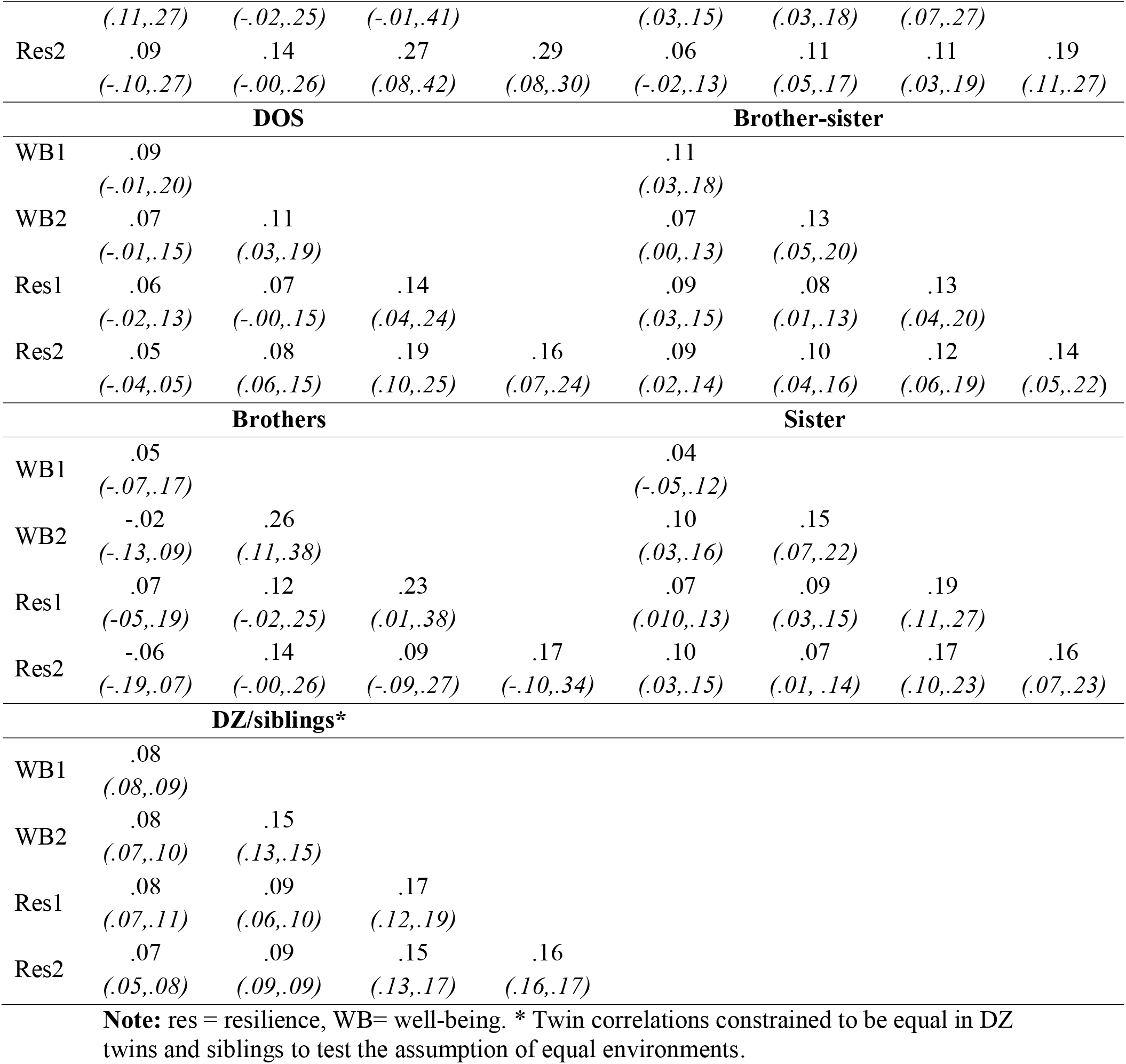
The twin and twin-sibling correlations for resilience and well-being within (diagonal) and cross traits and time points (off-diagonal).

The full bivariate ADE measurement model with sex differences is shown in supplementary Figure S4. First, we tested the quantitative sex effect by constraining all path estimates to be equal for males and females (see Table 4, model 2). This model gave a significant deterioration of fit (p<.001). Next, we tested if only the latent factor path estimates could be constrained to be equal in males and females, whereas the path estimates of the time-specific factors were allowed to differ. This model did not lead to a deterioration of the fit, p=.400. Next, both specific and common dominant genetic effects (D) did not contribute significantly to the (co)variance (p=.269). Therefore, the final model is an AE model without sex differences in the latent factor, but with sex differences in the time-specific factors (see Figure 4).

**Figure 4.**
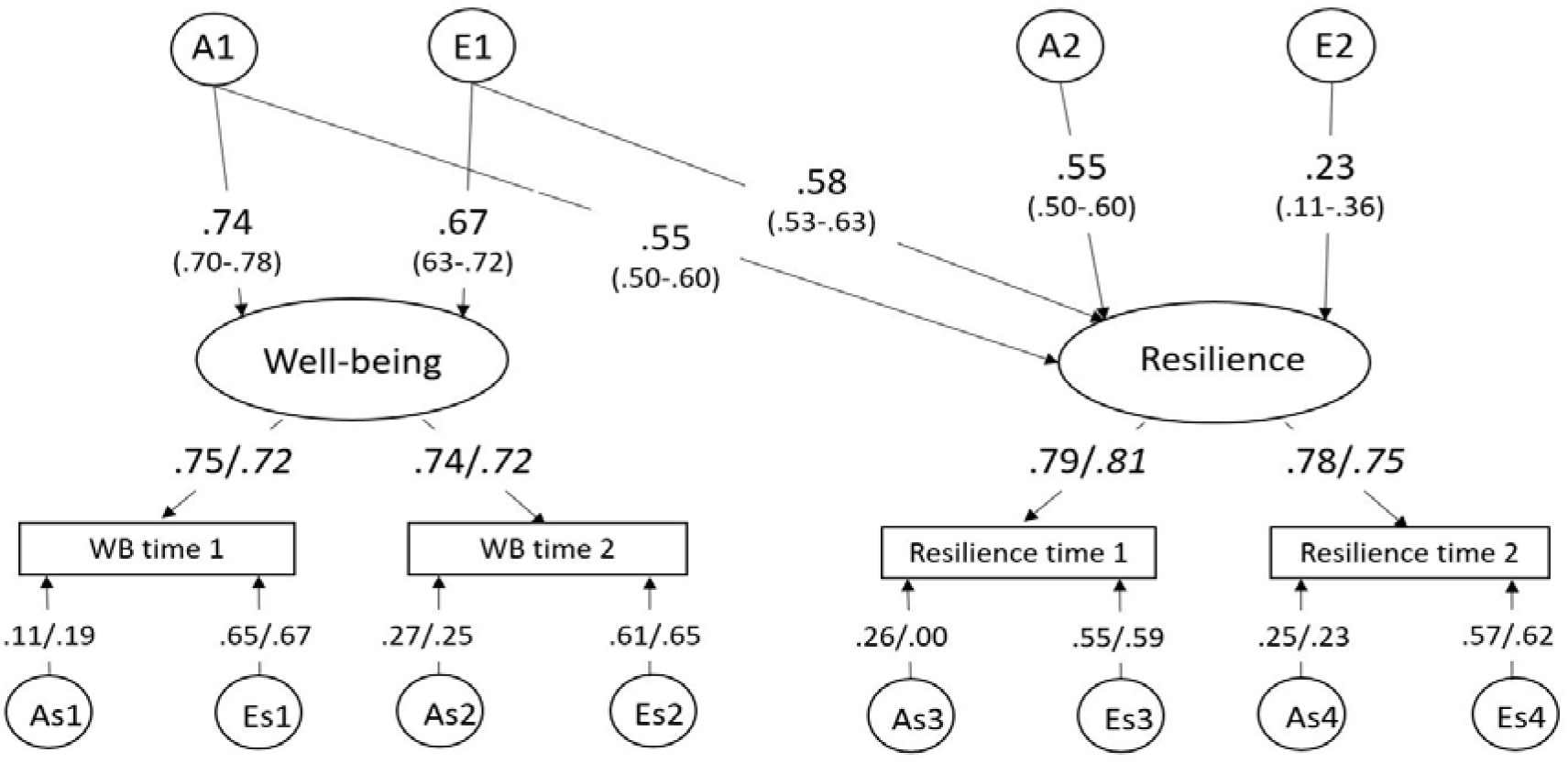
Unstandardized path estimates of the final common pathway model of resilience and well-being. The factor loadings from the common factors to the time-specific factors and the time-specific variance decomposition could not be constrained to be equal for females and males, indicated by estimates for females/males. WB= well-being, A= common additive genetic effect, E= common unique environmental effects, As= time-specific additive genetic effect, Es= time-specific environmental effect.

**Table 4.** Results of the model fitting for the psychometric model for resilience and well-being.

| Model | Variables | Constraints | vs | -2LL | df | AIC | $\Delta$ -2LL | $\Delta$ df | p |
| --- | --- | --- | --- | --- | --- | --- | --- | --- | --- |
| I | ADE |  |  | 176426.98 | 27034 | 122358.98 |  |  |  |
| II | ADE | Equal sex | I | 176614.80 | 27067 | 122480.80 | 187.82 | 33 | 1.43E-23 |
| III | ADE | Equal sex only<br>in latent part | I | 176436.40 | 27043 | 122350.40 | 9.42 | 9 | .400 |
| **IV** | **ADE** | **D=0** | **III** | **176449.79** | **27054** | **122341.79** | **13.39** | **11** | **.269** |

*Note:* df= degrees of freedom.

In the final bivariate model, the heritability of the latent well-being factor is estimated at 54.8% (95% CI: 53.1-57.1), whereas the unique environment explains 45.2% (43.1-51.4) of the variance in well-being. The heritability of the latent resilience factor is 60.9% (95% CI: 60.6-62.3) and the unique environment explains 39.1% (38.9-41.2) of the variance. At time 1 and time 2, time specific genetic influences explained respectively 32% and 37% of the variance in well-being in females. For males, the time specific heritability of well-being was similar, with 32% and 35% at T1 and T2 respectively. For resilience, the time specific heritability was 45% and 43% for females, but lower for males, with 39% and 36% at T1 and T2 respectively.

Of the covariance between resilience and well-being, 51.2% is explained by genetic factors and 48.8% by environmental factors. The genetic and environmental correlation between the latent factors of resilience and well-being are .71 (95% CI: .70-.71) and .93 (95% CI: .86-.98) respectively. As expected under the causal model, the genetic and environmental correlations could not be constrained to zero, p<.001 (see supplementary Table S3).

#### 3.4.3. Longitudinal twin-sibling models

In a bivariate longitudinal twin model with the resilience score at T1 and well-being at T2, we could not constrain the estimates to be equal across sex. Thus we tested the significance of the genetic and environmental correlations separately for males and females, by constraining the covariance between resilience and well-being. In line with the measurement model, we dropped D dropped (p=.005). The genetic correlations from resilience at baseline and well-being at T2 were .62 (95% CI: .53-.74) and .63 (95% CI:.41- .85) for females and males respectively. The environmental correlations were .19 (95% CI:.11-.26) and .23 (95% CI:.10-.35). Constraining any of the correlations to zero resulted in a deterioration in fit (p<.001) (see supplementary Table S4), in line with a causal relationship from resilience at T1 to well-being at T2.

In a bivariate model with the well-being score at T1 and resilience at T2 separately for males and females, we dropped D (p=.002). The genetic correlation from well-being at baseline and resilience a few years later were .64 (95% CI: .52-.76) and .29 (95% CI: .24-.55) for females and males respectively. The environmental correlations were .20 (95% CI: .12-.27) and .32 (95% CI: .19-.43). Constraining the genetic correlation to zero in females or the environmental correlation to zero in males and females resulted in a deterioration of the model fit (p<.001). In males, constraining the genetic correlation to zero did not change the model fit (p=.033) (see supplementary Table S5) which seems to falsify the causal hypothesis in males.

#### 3.4.4. MZ twin difference model

The MZ twin intrapair differences model showed that regressing the resilience MZ twin difference score on the well-being MZ twin difference score resulted in significant estimates at both time points (T1: β=.38, SE=0.03, R^2^ = 0.15, p<.001, T2: β=.47, SE=0.02, R^2^ = 0.21, p<.001). Similarly, regressing the well-being difference score of MZ pairs on the resilience difference score resulted in significant estimates (T1: β=.40, SE=0.03, R^2^ = 0.15, p<.001, T2: β=.44, SE=0.02, R^2^ = 0.21, p<.001) (see Figure 5 upper panels).

**Figure 5.**
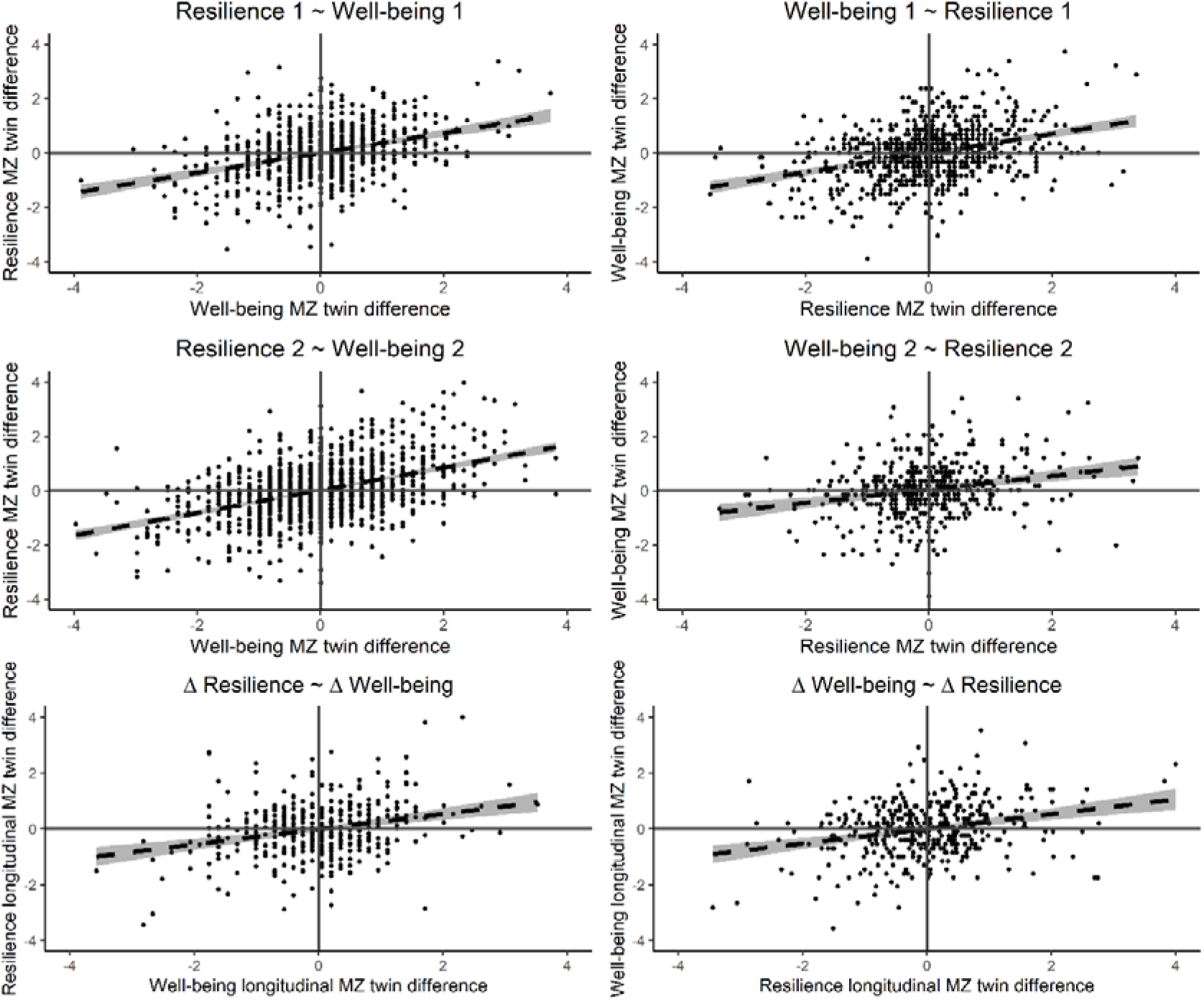
The monozygotic twin differences models. In the left upper panels, the MZ twin difference score of resilience is predicted by MZ twin differences in well-being (crosssectional, at T1 and T2). In the right upper panels, the MZ twin difference score of well-being is predicted by MZ twin differences in resilience (cross-sectional, at T1 and T2). The lower panel shows the longitudinal association between MZ within-twin pair differences in resilience and well-being.

The MZ twin longitudinal intrapair differences model showed a significant estimate from regressing the resilience change difference score on the well-being change difference, β=.32, SE=0.05, R^2^ = 0.10, p<.001. Similarly, regressing the well-being difference score on the resilience difference score resulted in a significant estimate, β=.33, SE=0.05, R^2^ = 0.10, p<.001 (see Figure 5 lower panel).

These findings are in line with a possible causal relation between resilience and well-being, indicating that higher well-being can lead to higher resilience and/or vice versa. As MZ twins share 100% of their genes, genetic confounding is taken into account.

#### 3.4.5. MR-DoC model

Lastly, we included the PGS as an instrumental variable in the twin model, applying the MR-DoC model (Minică et al., 2018) that can model causal effects while relaxing the MR no pleiotropy assumption. Due to data limitations (i.e. the resilience PGS is not powerful in predicting resilience), we could only test the effect of well-being on resilience. When including the PGS for well-being, estimating the pleiotropic effect freely and constraining the environmental correlation (rE) to zero, the direct effect from well-being to resilience cannot be dropped from this model at both times, p<.001 (see Figure 6 and supplementary Table S6 and S7), in line with a causal relation from well-being to resilience. The explained variance in resilience by well-being is 11.6% and 19.4% at time point 1 and 2 respectively. In addition, as expected, there is pleiotropy between well-being and resilience as indicated by the significant path b2 from the well-being PGS to resilience.

**Figure 6.**
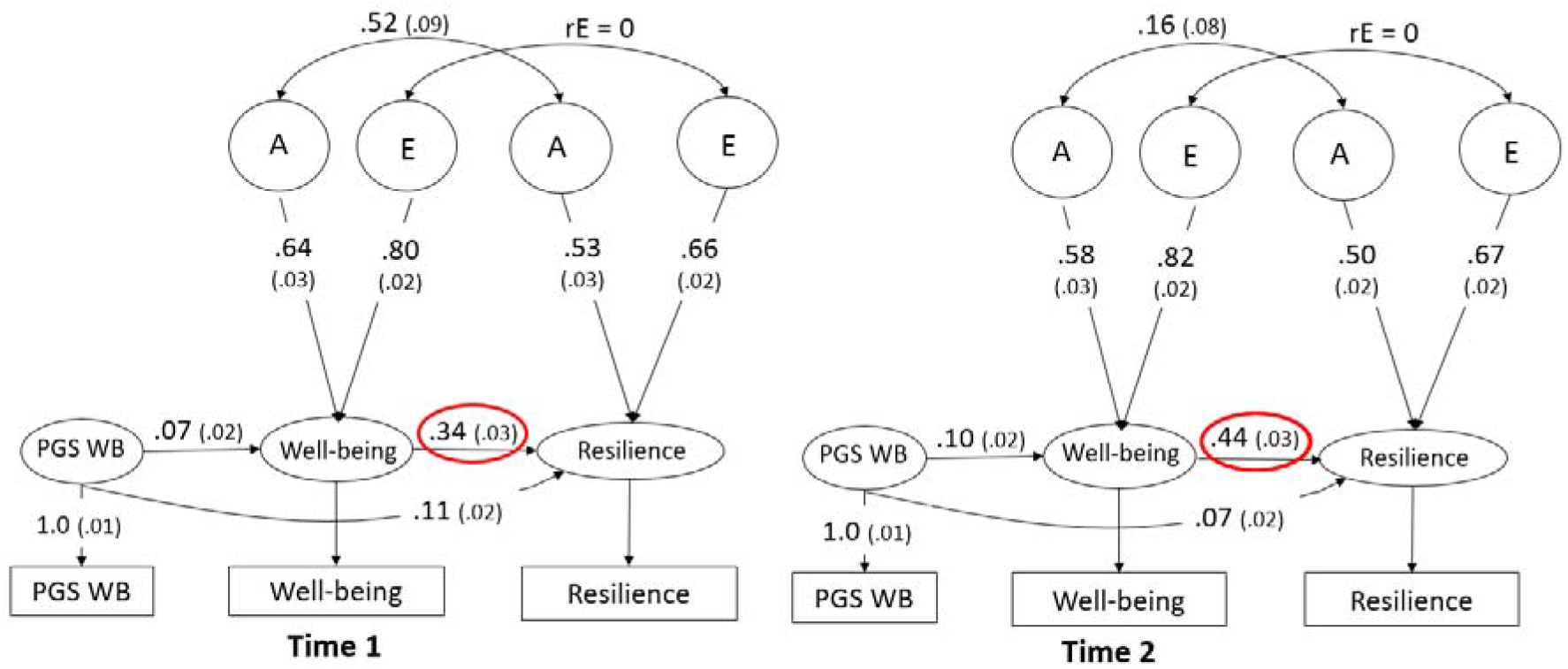
The results of the MR-DoC model. The models show the model with the wellbeing polygenic score and resilience as the outcome in cross-sectional data from time point 1 and 2 respectively. The causal effects can be seen in the red circle. WB= well-being,, PGS= polygenic score, A= additive genetic effect, E= unique environmental effect, rE= environmental correlation.

We cannot freely estimate the environmental correlation (rE) in the model, as both well-being resilience are AE traits. However, we can fix the rE to a correlation of various sizes instead of fixing the correlation to zero. As exploratory analyses we ran different MR-DoC models fixing the rE to respectively 0.1, 0.2, 0.3, 0.4, 0.5, and 0.8 at both time points (see supplementary Table S8 for the model fits). As indicated by the equal model fits, we did not have the power to distinguish between the model fit of a model with rE is 0,0.1, 0.2 and 0.3. At time point 1 (and similarly for time point 2), the estimate of the causal effect from well-being to resilience decreased from 11.6% when rE=0 to respectively 6.6%, 3.0% and 0.7% with a rE of 0.1, 0.2 and 0.3. With a fixed rE of 0.4 and higher, the model fit started to decrease and the causal effect estimate was not significant anymore.

## 4. Discussion

### 4.1. Summary of results

We investigated the association between resilience and well-being in a large sample of twins and their siblings from the Netherlands Twin Register and tested whether the observed overlap between resilience and well-being was due to bidirectional causal effects, after taking into account genetic possible genetic overlap between the two traits. The twin-sibling models showed strong cross-sectional and longitudinal correlations and a large overlap in genetic and environmental factors underlying resilience and well-being. There was a sex effect in the resilience mean score (men showed more resilience than women), but no sex difference in the genetic architecture of the latent factors for resilience and well-being. Results based on studies using GWAS summary statistics provided weak support for causal effects. Polygenic score analyses showed that the genetic risk for well-being is a predictor for resilience, but the genetic risk for self-reported resilience did not predict indirect resilience or well-being. The results of different causality analyses in twin-sibling models (De Moor et al., 2008) were not successful in falsifying the bidirectional causal relation between resilience and well-being. The explicit and most informative test of causality, the MR-DoC model using both twin and PGS data, supported the unidirectional causal hypothesis from well-being to resilience, whereas we could not test the causal hypothesis from resilience to well-being, due to power issues in the resilience GWAS data.

Using a psychometric twin model with data of two time points, the heritability estimates for the latent traits well-being and resilience were similar with respectively 54.8% and 60.9%. More than half of the variance in the stable part of resilience and well-being is explained by genetic factors. Whereas the heritability estimates at the two time points were around 30-40% for both traits, the heritability estimates of the psychometric model were higher, as this is not confounded by measurement error and time-specific influences. These estimates are highly comparable to earlier studies on resilience and well-being (Amstadter et al., 2014; Bartels, 2015; Boardman et al., 2008).

About 50% of the covariance between resilience and well-being is explained by genetic factors and the strong genetic correlation (.71) indicates that the genetic factors underlying resilience and well-being overlap significantly. Environmental factors explain the other half of the covariance between resilience and well-being. The environmental correlation between the latent traits is close to unity (.93) and indicates an almost perfect overlap in the environmental factors influencing both traits.

The various analyses using genetically informative samples were not successful in falsifying the causal hypothesis, even when correcting for genetic confounding. Therefore, we suggest that our findings are in line with a bidirectional causal relation between resilience and well-being instead of an underlying set of genes and/or environmental factors. To strengthen this finding, we applied the MR-DoC model. The MR-DoC model allows us to estimate causal effects, even in the presence of pleiotropy between the phenotypes. This model yielded results consistent with a causal relation from well-being to resilience, with about 11% (T1) and 20% (T2) of the variance in resilience explained by a causal effect from well-being. Due to the limited power of the resilience PGS, the causality in the other direction, from resilience to well-being could not be tested reliably in the MR-DoC model.

The assumption that the unique environmental sources of variation for resilience and well-being equals zero is necessary in the MR-DoC model for identification and this might seem implausible as we find an environmental correlation of .93 for well-being and resilience in the latent twin-sibling model. However, the unique environmental effects on well-being can be related with resilience via the causal path only, where the unique environmental effects influence the outcome via its effect on the exposure and not directly. In bivariate twin models, this causal path is missing, what could result in a large environmental correlation.

As exploratory analyses, we fixed the environmental correlation to various sizes instead of fixing the correlation to zero. This led to equal model fits for an environmental correlation of 0, 0.1, 0.2 and 0.3, as we did not have the power to distinguish between these models. At time point 1, the estimate of the causal effect from well-being to resilience decreased from 11.6% when the environmental is zero to respectively 6.6%, 3.0% and 0.7% with a rE of 0.1, 0.2 and 0.3. With a higher environmental correlation, the model fit started to decrease and the causal effect estimate was not significant anymore. These results indicate that the model does find a causal effect between well-being and resilience. However, it is likely that there is an environmental correlation between well-being and resilience as well, reducing the size of the causal effect. More power is needed to determine the size of the causal effect.

Another explanation for the strong correlations between resilience and well-being could be a third variable underlying both traits and explaining the bidirectional relationship between them. For example, self-rated general health has a strong genetic correlation with well-being (Baselmans, van de Weijer, et al., 2019), although the direction of causation between well-being and health is not clear (Rohrer & Lucas, 2020). If general health (or another variable) causally influences both well-being and resilience, a strong correlation between well-being and resilience does not necessarily mean the constructs have an influence on each other. Future research should include such variables in one analysis to investigate this possibility. For now, based on the converging results of the different analyses, we suggest resilience and well-being might have some causal effects on each other.

### 4.2. Points of discussion and limitations

#### 4.2.1. Defining resilience and well-being

There is discussion about definition of resilience and well-being resulting in no universal or commonly agreed upon definition. Different questionnaires are validated to assess self-report resilience, like the Ego-Resilience scale (Block & Kremen, 1996) and Connor-Davidson Resilience Scale (Connor & Davidson, 2003). More recently, researchers are emphasizing the need for improved operationalizations of resilience (Kalisch et al., 2019; Stainton et al., 2019). Resilience is not a stable trait but a complex, interactive process leading to positive psychological outcomes in response to stress or adversity (Kalisch et al., 2017). In line with these definitions, in our sample, direct self-reported resilience seems to be different from resilience measured as the response to exposure to stress, based on the nonsignificant polygenic score predictions in our study. Therefore, the results also underscore the need for a clear and commonly agreed upon definition of resilience.

We defined resilience as the better than predicted psychological outcome based on the number of stressful life events experienced. A difficulty in this definition is the inclusion of the type and number of life events experienced. In our study, we included 16 and 19 (time 1 and 2) life events about illness, dead of close others and events like robbery and accidents. This is not an exhaustive list of life events and the personal impact of life events might differ per individual. Therefore, further research should weight the personal impact of life events to better operationalize resilience.

Furthermore, whereas we focused on the absence of psychopathological symptoms after stress as our resilience measure to compare the overlap with well-being, another approach to measure resilience is to assess the positive adaptation after stress. As mental health is more than the absence of psychopathology, instead of assessing anxious-depressive symptoms, well-being might be assessed as the outcome measure after stress. If people experience higher well-being than expected based on the stress experienced, this might be a sign of resilience as well.

Multiple theories about the definition of well-being exist as well and as mentioned in the introduction, often a distinction between subjective and psychological well-being is made. However, subjective and psychological well-being are strongly related, both genetically and phenotypically (e.g. Baselmans et al., 2019; Joshanloo, 2016). We included life satisfaction as a measure of well-being, but we expect a similar overlap between other aspects of wellbeing and resilience, because of the strong overlap between the different well-being measures.

#### 4.2.2. Resilience GWAS

The GWAS summary statistics used to compute polygenic scores for resilience were based on the only GWAS to date on (self-reported) resilience (Stein et al., 2019). The GWAS in the relatively small US army soldiers sample (N=11.492) resulted in one independent significant locus. The use of these GWAs summary statistics comes with the following limitations. First, the discovery sample size is small, resulting in less power to detect genetic associations and subsequently less power to predict resilience using the summary statistics of the GWAS (Dudbridge, 2013). Secondly, a sample restricted to soldiers might not reflect the general population (i.e., results based on this sample might not generalize to the population). Third, in contrast to our indirect measure of resilience, the GWAS included a direct measure of self-reported resilience (STARRS 5-item questionnaire, rating of the ability to handle stress). Therefore, the PGS based on these summary statistics reflects the genetic risk for selfreport resilience.

As can be seen in our results, the power to detect association between the resilience PGS and the resilience measure was low. The resilience PGS did predict indirect resilience to some extent, but the variance explained was almost zero (<.001%). Furthermore, although there is an indication for a genetic relation and overlap between resilience and well-being, the self-reported resilience PGS did not predict well-being. Due to the low power of the resilience PGS and the small association between the resilience PGS and resilience score, applying the MR-DoC model including the resilience PGS would not lead to reliable results. Not all MR assumptions (i.e. the strong association between the genetic instrument and exposure) are fully met.

The MR-DoC model can be extended to test a bidirectional relationship, including both PGS of resilience and well-being at the same time. Such a model can strengthen the results and the constraint of the environmental correlation to zero is not necessary anymore. However, for this model two sets of powerful polygenic scores are needed. As the resilience PGS lacks power, we did not model such a bidirectional MR-DoC model. Similarly, as the resilience GWAS is not predictive of our outcome-based measure of resilience and did not have much power, we did not apply SNP based MR methods, such as two-sample MR methods, like MR-Egger. Even if these methods would show causality, the results will not be informative about the relation between well-being and outcome-based resilience, as it has been shown that there is only a moderate degree of genetic overlap between self-reported and outcome-based resilience, using twin models (Sawyers et al., 2020).

The less powerful PGS for resilience due to the small GWAS discovery sample and different operationalization of resilience limits the interpretation of the molecular genetic analyses in our study. To replicate and strengthen our findings on the overlap and direction of effect between resilience and well-being with molecular genetic data, a powerful GWAS for resilience as response to stress (instead of direct self-reported resilience) carried out in a large sample from the general population is needed. In line with our operationalization, a measure of internalizing problems (e.g. anxiety and/or depression) and the number of experienced life events can be combined to create such a measure of resilience in a large genotyped sample.

Lastly, as with most GWASs, we used the summary statistics resulting from studies restricted to individuals with European ancestry. Therefore, our results might not generalize to populations of different genetic ancestries. Recently, large projects to include individuals from other ancestries have been started as analyzing a more inclusive and diverse dataset might increase power to detect associations (Pan-UKB-team, 2020).

### 4.3. Implications

The results in our large genetically informative sample suggest a large overlap and a potential bidirectional relationship between resilience (psychological outcome after negative life events) and well-being (life satisfaction). If replicated, the results implicate that increasing well-being might lead to increased resilience (i.e. a positive psychological outcome after negative life events or trauma) as well and vice versa. As resilience and well-being are both negatively related to psychopathology (Amstadter et al., 2016; Diener et al., 2017; Greenspoon & Saklofske, 2001; Howell et al., 2007), the bi-directionality between the positive constructs of resilience and well-being can have implications for interventions to prevent or lower vulnerability for psychopathology. Increasing well-being can be important to prevent trauma-related psychopathology and psychiatric symptoms. Vice versa, increasing resilience (i.e. decreasing the likelihood of psychopathological or psychiatric symptoms after trauma) can protect an individual’s well-being after stress. The independent interventions related to increasing well-being and separate interventions for coping with trauma might supplement each other.

## Acknowledgements

The authors thank all NTR participants, who participated in this study. In addition, we would like to thank Wouter Peyrot for valuable feedback on an earlier version of this manuscript.

## Supplementary tables and figures

**Table S1.** Number of participants per zygosity and sex for every variable.

|  | Resilience 1 |  |  |  | Well-being 1 |  |  |  | Resilience 2 |  |  |  | Well-being 2 |  |  |  |
| --- | --- | --- | --- | --- | --- | --- | --- | --- | --- | --- | --- | --- | --- | --- | --- | --- |
|  | T 1 | T 2 | S m | S f | T 1 | T 2 | S m | S f | T 1 | T 2 | S m | S f | T 1 | T 2 | S m | S f |
| MZF | 769 | 717 | 127 | 187 | 795 | 747 | 126 | 198 | 1367 | 1259 | 144 | 220 | 1674 | 1508 | 171 | 267 |
| MZM | 323 | 264 | 74 | 71 | 333 | 274 | 76 | 74 | 506 | 441 | 69 | 99 | 649 | 551 | 87 | 121 |
| DZF | 412 | 336 | 67 | 86 | 433 | 354 | 70 | 92 | 708 | 574 | 82 | 124 | 891 | 709 | 96 | 142 |
| DZM | 178 | 128 | 35 | 46 | 184 | 125 | 35 | 47 | 293 | 224 | 40 | 66 | 394 | 283 | 49 | 78 |
| DZO | 392 | 358 | 82 | 110 | 405 | 377 | 83 | 119 | 629 | 621 | 89 | 147 | 914 | 813 | 105 | 183 |
| Total | 2074 | 1803 | 385 | 500 | 2150 | 1877 | 390 | 530 | 3503 | 3119 | 424 | 656 | 4522 | 3864 | 508 | 791 |

*Note:* T1= twin 1, T2= twin 2, Sm= sibling male, Sf= sibling female.

**Table S2.** The stressful life events included in the questionnaire at both time points.

| Time 1 | Time 2 |
| --- | --- |
| 1. Death of life partner, | 1. Financial problems, |
| 2. Death of father, | 2. Job loss, |
| 3. Death of mother, | 3. Drop out of education, |
| 4. Death of child, | 4. Relationship problems with partner, |
| 5. Death of sibling, | 5. Relationship problems with child, |
| 6. Death of other loved one, | 6. Relationship problems with other loved one, |
| 7. Serious disease of yourself, | 7. Getting hospitalized, |
| 8. Serious disease of life partner, | 8. Serious illness yourself, |
| 9. Serious disease of child, | 9. Serious illness partner, |
| 10. Serious disease of other loved one, | 10. Serious illness child, |
| 11. End of relation, | 11. Serious illness parent, |
| 12. Traffic accident, | 12. Serious illness other loved one, |
| 13. Violent crime, | 13. Death of partner, |
| 14. Sexual crime, | 14. Death of child, |
| 15. Theft, | 15. Death of other loved one, |
| 16. Getting fired. | 16. Traffic accident, |
|  | 17. Theft, |
|  | 18. Violent crime, |
|  | 19. Sexual crime. |

**Table S3.** Results of the psychometric model constraining the genetic and environmental correlation.

| Base | Comparison | -2LL | df | AIC | ΔLL | $\Delta$ df | p |
| --- | --- | --- | --- | --- | --- | --- | --- |
| **AE** |  | 176449.79 | 27054 | 122341.79 |  |  |  |
| AE | rA=0 | 176731.08 | 27055 | 122621.08 | 281.29 | 1 | 3.93E-63 |
| AE | rE=0 | 176966.24 | 27055 | 122856.24 | 516.46 | 1 | 2.50E-114 |

*Note:* rA= genetic correlation, rE= environmental correlation.

**Table S4.** Results of the longitudinal twin model fitting for resilience at baseline and well-being a few years later.

| Base | Comparison | ep | -2LL | df | AIC | ΔLL | Δdf | p |
| --- | --- | --- | --- | --- | --- | --- | --- | --- |
| **AE** |  | 16 | 93841.40 | 14431 | 64979.40 |  |  |  |
| AE | rAf=0 | 15 | 93946.52 | 14432 | 65082.52 | 105.13 | 1 | <.0001 |
| AE | rAm=0 | 15 | 93867.60 | 14432 | 65003.60 | 26.20 | 1 | <.0001 |
| AE | rEf=0 | 15 | 93863.40 | 14432 | 64999.37 | 21.97 | 1 | <.0001 |
| AE | rEm=0 | 15 | 93853.11 | 14432 | 64989.12 | 11.73 | 1 | <.0001 |

*Note:* ep= estimated parameters, rAf= genetic correlation females, rAm= genetic correlation males, rEf= environmental correlation females, rEm= environmental correlation males.

**Table S5.** Results of the longitudinal twin model fitting for well-being score at baseline and resilience a few years later.

| base | comparison | ep | -2LL | df | AIC | ΔLL | Δdf | p |
| --- | --- | --- | --- | --- | --- | --- | --- | --- |
| AE |  | 16 | 86832.88 | 12633 | 61566.88 |  |  |  |
| AE | rAf=0 | 15 | 86909.84 | 12634 | 61641.84 | 76.96 | 1 | <.0001 |
| AE | **rAm=0** | 15 | 86837.44 | 12634 | 61569.44 | 4.55 | 1 | **.0329** |
| AE | rEf=0 | 15 | 86855.64 | 12634 | 61587.64 | 22.76 | 1 | <.0001 |
| AE | rEm=0 | 15 | 86854.52 | 12634 | 61586.52 | 21.63 | 1 | <.0001 |

*Note:* ep= estimated parameters, rAf= genetic correlation females, rAm= genetic correlation males, rEf= environmental correlation females, rEm= environmental correlation males.

**Table S6.** Results of the MR-DoC model fitting for the different time points.

| base | comparison | ep | -2LL | df | AIC | ΔLL | Δdf | p |
| --- | --- | --- | --- | --- | --- | --- | --- | --- |
| **Time 1** |  |  |  |  |  |  |  |  |
| WB -> RES |  | 35 | 22796.60 | 8442 | 5912.6 |  |  |  |
| WB -> RES | g1=0 | 34 | 22898.07 | 8443 | 6012.1 | 101.5 | 1 | 7.27E-24 |
| **Time 2** |  |  |  |  |  |  |  |  |
| WB -> RES |  | 35 | 30263.71 | 11243 | 7777.707 |  |  |  |
| WB -> RES | g1=0 | 34 | 30560.39 | 11244 | 8072.391 | 296.684 | 1 | 1.74E-66 |

Note: WB= well-being, RES= resilience, g1= causal effect, ep= estimated parameters, df= degrees of freedom.

**Table S7.** Estimates of the MR-DoC models for the different time points.

| Standardized | name | Time 1:<br>WB -> Resilience |  | Time 2:<br>WB -> Resilience |  |
| --- | --- | --- | --- | --- | --- |
|  |  | Estimate | Std.Error | Estimate | Std.Error |
| Causal effect | g1 | 0.337 | 0.033 | 0.443 | 0.025 |
| PGS exposure | b1 | 0.077 | 0.022 | 0.098 | 0.017 |
| PGS outcome | b2 | 0.109 | 0.019 | 0.069 | 0.016 |
| A effect exposure | ab | 0.638 | 0.032 | 0.581 | 0.025 |
| E effect exposure | eb | 0.795 | 0.022 | 0.816 | 0.016 |
| A effect outcome | as | 0.530 | 0.029 | 0.502 | 0.022 |
| E effect outcome | es | 0.659 | 0.018 | 0.673 | 0.015 |
| PGS to PGS | x | 1.001 | 0.012 | 1.001 | 0.012 |
| Genetic correlation | ra | 0.521 | 0.090 | 0.159 | 0.082 |

Note: WB= well-being.

**Table S8.** Model fit of models with various fixed rE values.

| **Time 1** | rE | parameters | -2LL | df | AIC | g1 | R2 (%) |
| --- | --- | --- | --- | --- | --- | --- | --- |
|  | 0 | 35 | 8442 | 22796.6 | 5912.6 | 0.337 | 11.36% |
|  | 0.1 | 35 | 8442 | 22796.6 | 5912.6 | 0.2536 | 6.43% |
|  | 0.2 | 35 | 8442 | 22796.6 | 5912.6 | 0.168 | 2.81% |
|  | 0.3 | 35 | 8442 | 22796.6 | 5912.6 | 0.076 | 0.58% |
|  | 0.4 | 35 | 8442 | 22797.1 | 5913.08 | 0 | 0.00% |
|  | 0.5 | 35 | 8442 | 22811.3 | 5927.3 | 0 | 0.00% |
|  | 0.8 | 35 | 8442 | 23106.7 | 6222.7 | 0 | 0.00% |
| **Time 2** | rE | parameters | df | -2LL | AIC | g1 | R2 (%) |
|  | 0 | 35 | 11243 | 30263.7 | 7777.71 | 0.443 | 19.62% |
|  | 0.1 | 35 | 11243 | 30263.7 | 7777.71 | 0.36 | 12.96% |
|  | 0.2 | 35 | 11243 | 30263.7 | 7777.71 | 0.275 | 7.56% |
|  | 0.3 | 35 | 11243 | 30263.7 | 7777.71 | 0.184 | 3.39% |
|  | 0.4 | 35 | 11243 | 30263.7 | 7777.71 | 0.083 | 0.69% |
|  | 0.5 | 35 | 11243 | 30265.1 | 7779.07 | 1.1E-14 | 0.00% |
|  | 0.8 | 35 | 11243 | 30832.1 | 8346.13 | 3.9E-10 | 0.00% |

**Figure S1.**
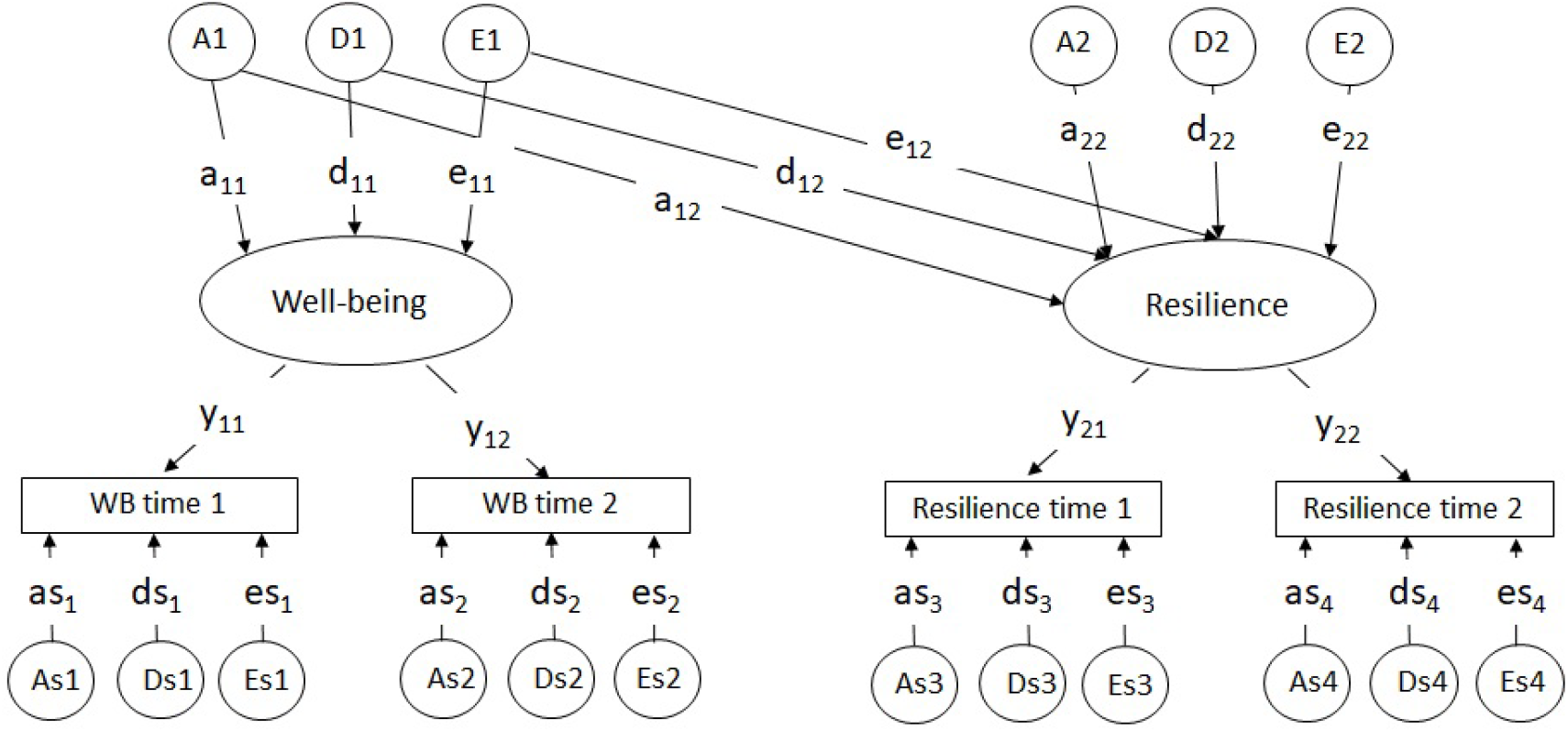
Longitudinal measurement model. WB= well-being, A= common additive genetic effect, E= common unique environmental effects, As= time-specific additive genetic effect, Es= time-specific environmental effect.

**Figure S2.**
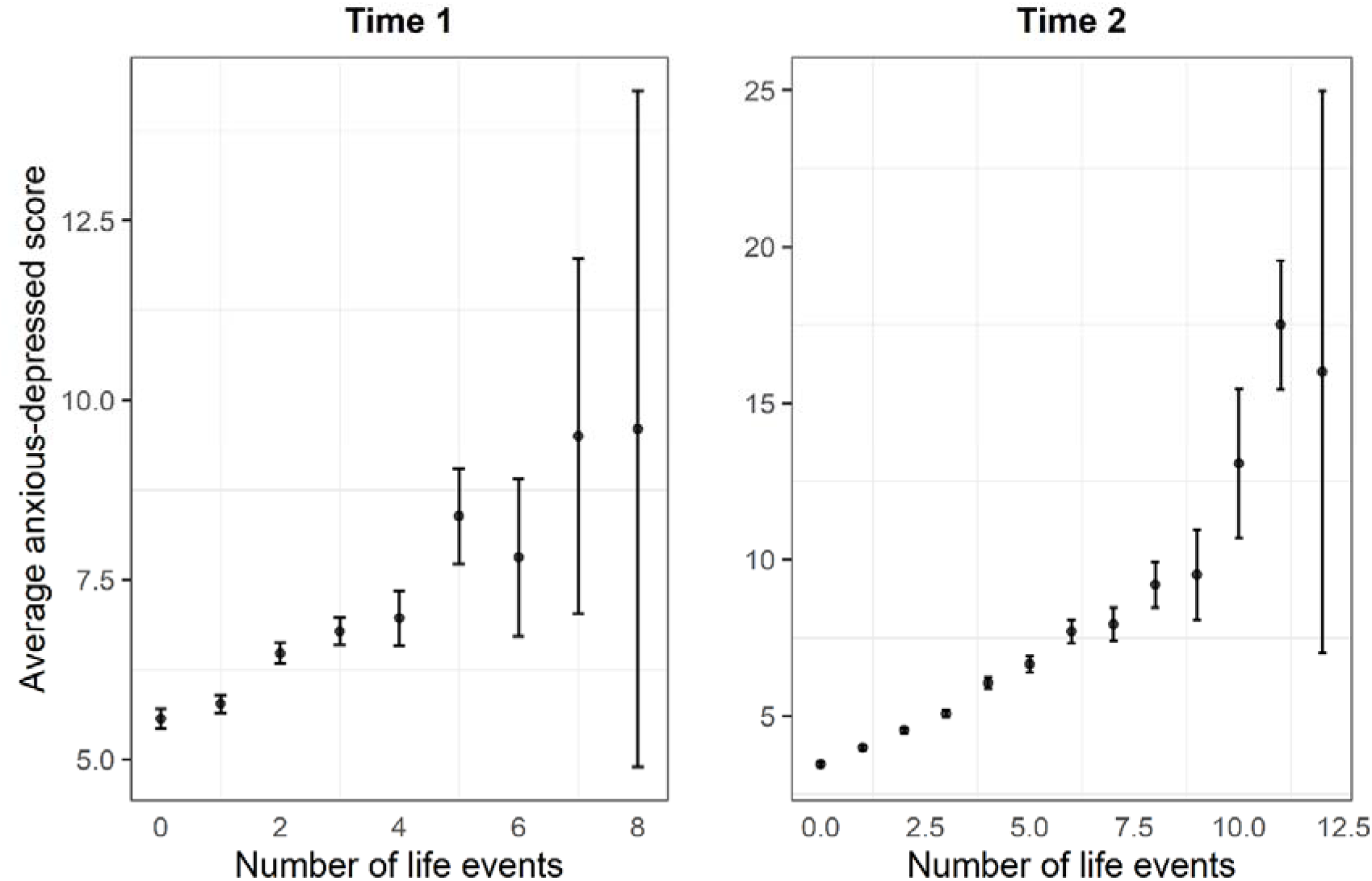
Association between the number of life events and the anxious depressed score.

**Figure S3.**
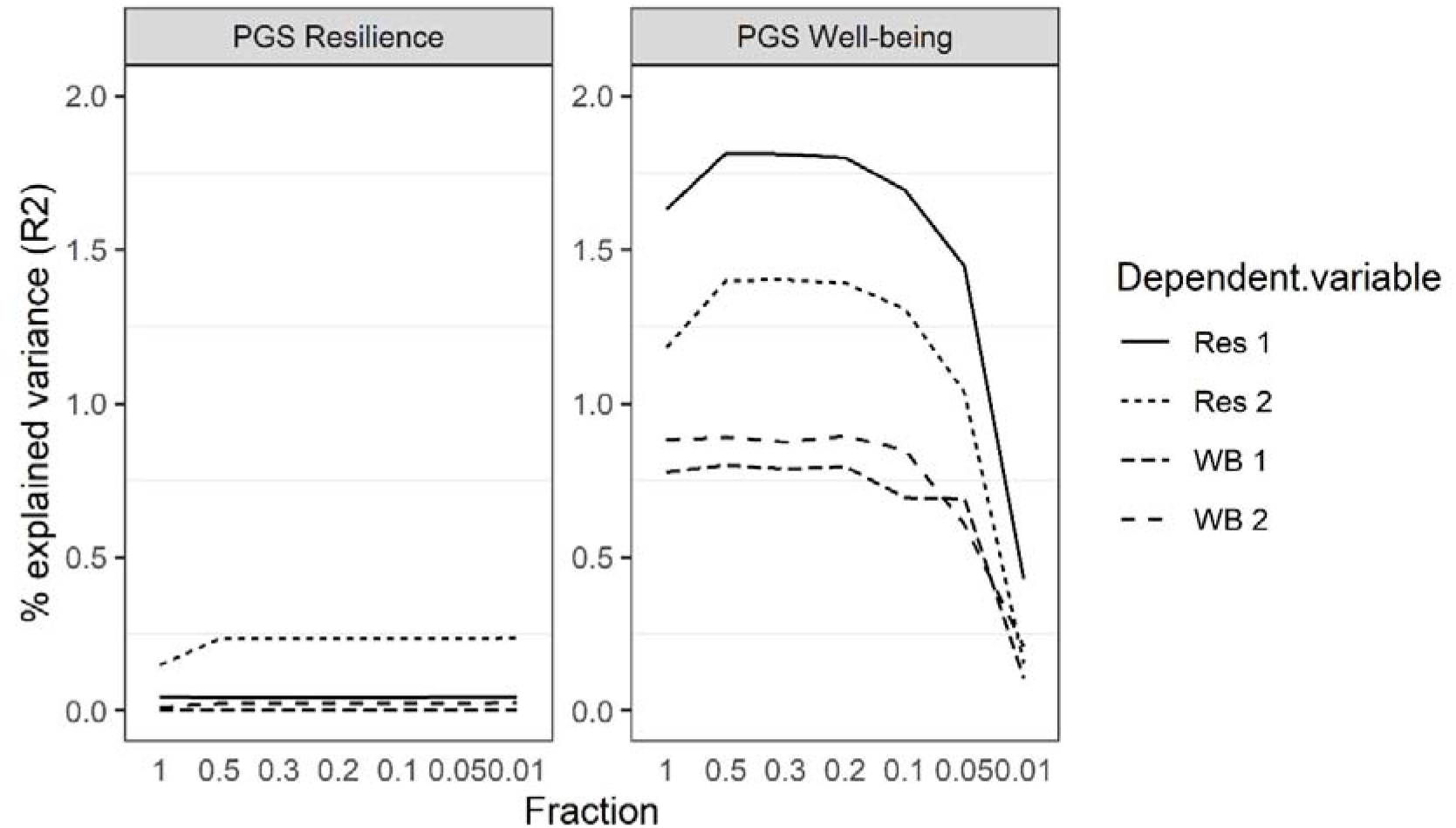
Explained variance by the resilience and well-being PGS for the different fractions of SNPs included.

**Figure S4.**
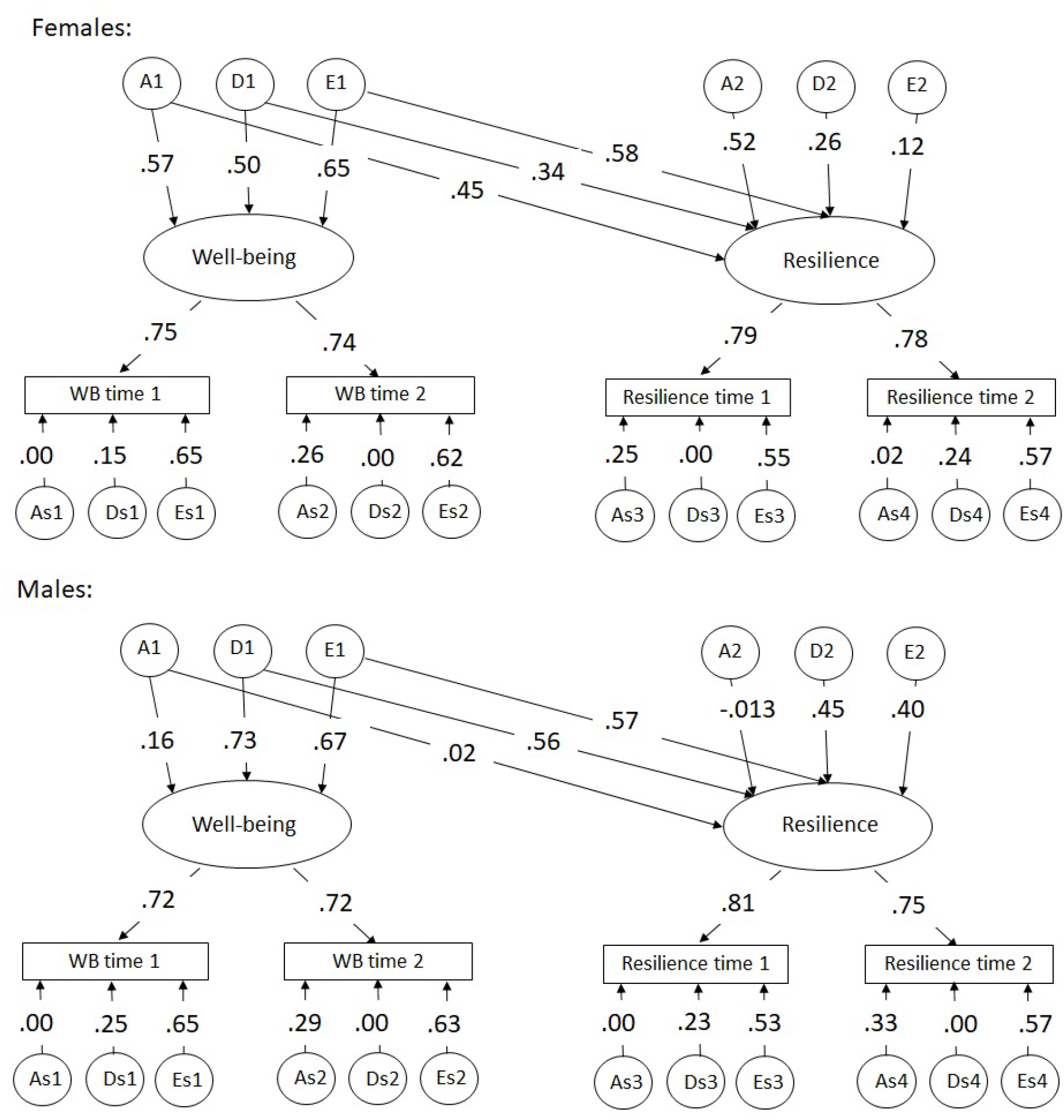
Path estimates of the full common pathway model of resilience and well-being for females and males. WB= well-being, A= common additive genetic effect, E= common unique environmental effects, As= time-specific additive genetic effect, Es= time-specific environmental effect.

